# Structural insights into Frizzled assembly by acylated Wnt and Frizzled Connector domain

**DOI:** 10.1101/2022.09.29.510206

**Authors:** Tao-Hsin Chang, Fu-Lien Hsieh, Karl Harlos, E. Yvonne Jones

## Abstract

Frizzled (Fz1-10) serve as the principal cell surface receptors for Wnt signalling. Aberrant expression of Fz is associated with cancer and neurodegeneration. The N-terminal extracellular domains of Fz include Cysteine-Rich Domain (CRD), Connector, and Linker. How the palmitoleate moiety (PAM) modified Wnt bound to Fz transduces the extracellular signal across the membrane remains incomplete. Here, we report the structures of Fz4 CRD and Connector (Fz4_CRD-Connector_), in complex with PAM modified Wnt7a peptide (PAM peptide). Fz4_CRD-Connector_ structures reveal an open-form of dimer – a flat-shaped hydrophobic groove to accommodate one PAM peptide at the dimer interface. Interestingly, the structure of Fz7_CRD_ bound to PAM peptide shows a dimeric closed-form – a curved-shaped hydrophobic groove at the dimer interface for one PAM peptide bound. We also reveal that Fz4_Connector_ has extensive interactions with Fz4_CRD_ and contributes to the Fz4 function. The studies shed insight on the development of novel strategies to modulate Fz function.

## Introduction

The wingless/int1 (Wnt) signalling pathway plays pivotal roles in cell-fate determination, embryonic development and adult tissue homeostasis^1–3^. Dysfunction of Wnt signalling leads to oncogenesis and degeneration diseases^4^. Thus, Wnt signalling is an attractive therapeutic target and many encouraging approaches have developed recently^1^. Mammalian genomes encode nineteen Wnt homologs that are secreted glycoproteins covalently linked with a palmitoleate moiety (PAM) – a 16-carbon *cis*-Δ9 unsaturated fatty acyl – at a conserved serine residue (e.g., Ser206 in human Wnt7a)^5,6^. The post-translational modification of PAM is critical for Wnt secretion^6^, Wnt binding to ten homologs of Frizzled receptors (Fz1-10)^7-9^, and Wnt signalling activity^5,10^, as well as Wnt movement forming the morphogen gradient^11^. The removal of PAM suppresses Wnt signalling via NOTUM, a Wnt specific deacylase^12,13^.

Fz is a multiple-domain protein which is composed of an N-terminal Cysteine-Rich Domain (CRD) followed by a Connector and a Linker to a seven-pass transmembrane domain and an intracellular domain (Fig.1). Fz is responsible to transduce Wnt signals across the plasma membrane, which is initially from acylated Wnt binding to Fz_CRD_^2,10^. Subsequently, the signals stimulate the Wnt intracellular signalosome formation^14,15^ and ultimately lead to activation of Wnt responsive gene transcription. Intriguingly, genetic functional studies further show that ten Fz homologs play extraordinarily diverse roles in developmental and homeostatic processes^2^ and these evolutionary close Fz homologs have a scenario of functional redundant and complement *in vivo^2^*. Furthermore, the protein levels of Fz receptors at the cell surface are stringently regulated by the complex of R-spondin, ZNRF3/RNF43 ubiquitin ligase, and LGR4/5/6 coreceptor to finetune Wnt signalling^16–19^.

**Figure 1.**
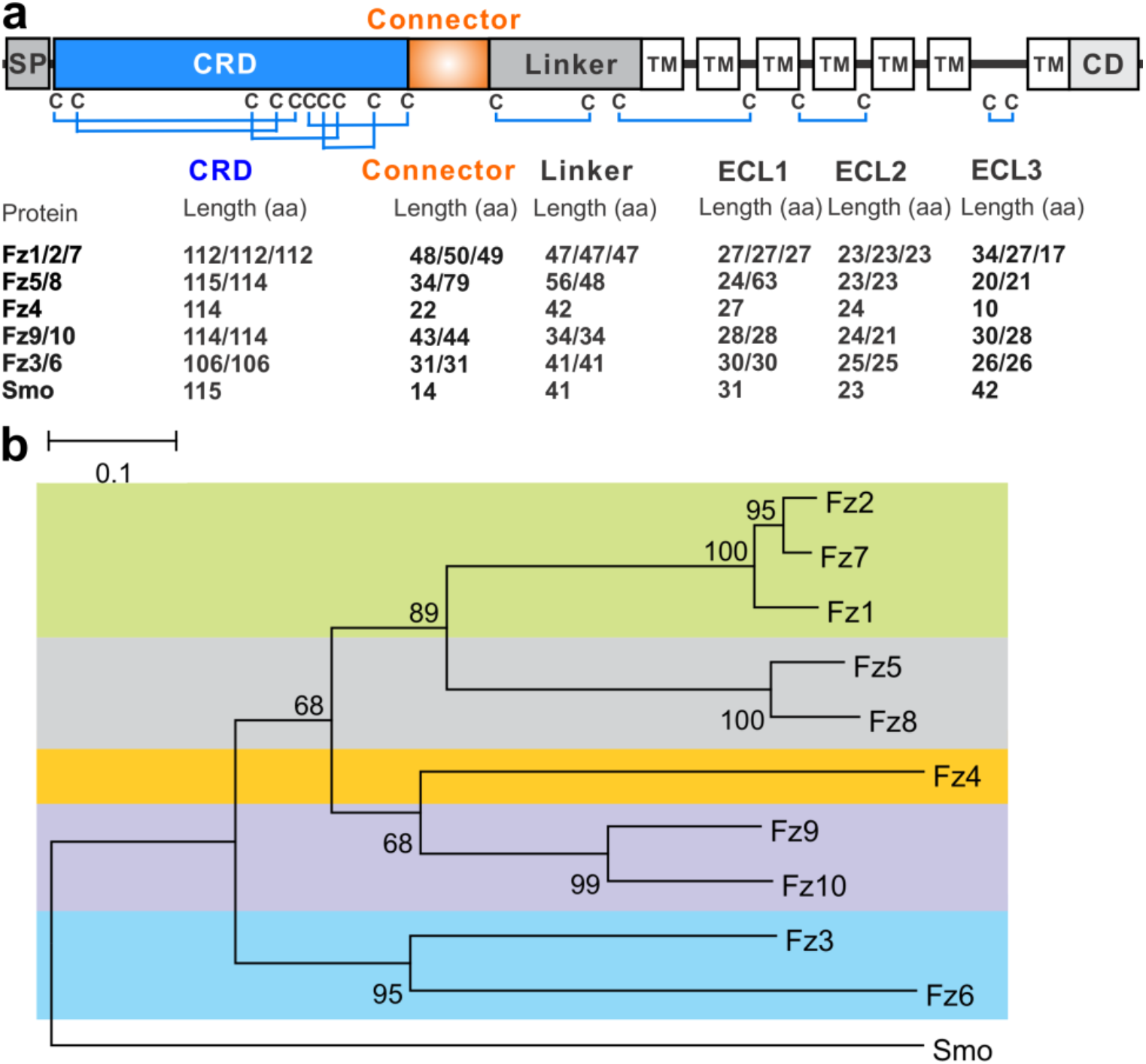
Domain architecture of Fz receptors and phylogenetic analysis of the segments containing CRD and Connector sequences. **(a)** Schematic representation of Fz receptors (SP, signal peptide; TM, transmembrane domain; CD, cytoplasmic domain). Conserved disulfide bridges among Fz receptors are drawn in blue lines. Extracellular segments of Fz receptors include a CRD, a Connector, a Linker, and three extracellular loops (ECL 1-3) between TMs. The protein lengths of those extracellular segments are listed below the diagram. **(b)** A protein sequence-based phylogenetic tree of the segment containing CRD and Connector sequences. Horizontal branch lengths correspond to evolutionary distance.

Long-standing questions in Wnt signalling include how nineteen Wnts pair with ten Fz receptors in an appropriate subset spatially and temporally and what the pair of Wnt and Fz studied *in vitro* can appropriately link to its *in vivo* biological functions. Previous functional studies had suggested both specificity and promiscuity occurred in Wnt and Fz pairs^2,7,20–22^. Furthermore, multiple lines of evidence from the structural studies have been brought into our understanding sharply, but partially. First, the structure of Xenopus Wnt8 (Wnt8) in complex with mouse Fz8_CRD_ reveals a 1:1 stoichiometry and two distinct binding sites^8^. In site 1, PAM on the N-terminal lobe of Wnt8 inserts into a hydrophobic groove of Fz8_CRD_ that is conserved across all Fz receptors. In site 2, a protruded hairpin loop on the C-terminal lobe of Wnt8 binds to a hydrophobic pocket of Fz8_CRD_. Much of these contact residues on Wnts and Fz receptors are evolutionarily conserved. Thus, how Wnt and Fz pairs achieve specificity and selectivity remains obscure. Second, prior studies used the chemical molecule of PAM that mimics the PAM on Wnt to co-crystalize with Fz4_CRD_ and Fz5_CRD_^23,24^. Their structural analyses revealed that the PAM locates at the hydrophobic dimer interface and acts like a hook to facilitate the dimerization^23,24^. Interestingly, the structure of Fz4_CRD_:PAM revealed an open-form dimer in contrast to a closed-form dimer of the Fz5_CRD_:PAM structure. Third, a recent structural study of human Wnt3 in complex with mouse Fz8_CRD_ revealed the same two binding sites as what was observed in Wnt8:Fz8_CRD_ complex but surprisingly presented a 2:2 stoichiometry where two PAMs on Wnts accommodate at the dimer interface^25^. Taken together, these studies raise interesting questions: how does the PAM on Wnt mediate different status of Fz (e.g. dimerization/oligomerization)? how does the 1:1 stoichiometry of Wnt and Fz transit into the 2:2 stoichiometry, further into the signalosome formation?

Here, we used (1) a disulfide bond cyclized peptide which corresponds to human Wnt7a (Cys202-Cys209) and attach with the PAM (refer to PAM peptide) to mimic the PAM on Wnt, rather than the chemical molecule of PAM used in previous studies^23,24^, and (2) Fz4 segments of CRD and Connector domains (Fz4_CRD-Connector_), and Fz7_CRD_ for co-crystallization with PAM peptide. The structures of Fz4_CRD-Connector_ in complex with PAM peptide at two crystal forms and Fz7_CRD_ in complex with PAM peptide were determined. The structural analyses revealed that (1) PAM peptide mediated the dimerization, (2) the structures of Fz4_CRD-Connector_:PAM peptide shows an open-form of dimeric arrangement, (3) the structure of Fz7_CRD_:PAM peptide shows a closed-form of the dimer, and (4) the PAM peptides present dynamic conformations while the overall positions are partially conserved in open-and closed-forms. More importantly, we revealed the segment structure of Fz4_Connector_ where the Connector has extensive interactions with the CRD. We also suggest that Fz4_Connector_ contribute to the Fz4 function, supported by the evidence of disease-associated mutations in Fz4_Connector_ and the functional assays of swapping the segment of Fz_Connector_. Taken together, our results provide insights into the roles of PAM on Wnts and Fz_Connector_ in Wnt signalling regulation and aid in the development of novel strategies to finetune Wnt signalling.

## Result

### Domain architecture and phylogenetic analysis of Fz receptors

We began our studies by analyzing the domain architecture of Fz receptors (Fig. 1a), while inspired by the full-length structure of a Fz evolutionarily related receptor, Smoothened^26–28^. We identified two additional segments that existed in the N-terminal Fz extracellular domain – Connector and Linker (Fig. 1a), except the CRD, the well-known segment of Fz receptors^2,9,10^. Specifically, CRD is generally around 106-112 amino acids defined from the first cystine to the tenth cystine that forms five disulfide bonds. CRD is the best-characterized domain for Wnt recognition^8–10,25^. This new annotated segment – Connector – starts after the tenth cystine of CRD and ends before the first cystine of Linker (Fig. 1a). More importantly, the Connector of Fz receptors reveals the semi-conserved in sequence length and amino acid composition in contrast to the CRD (Supplementary Fig. 1) and is a proline- and glycine-rich region. Furthermore, Linker – starting with the first cystine after Connector and ending before the first transmembrane domain of Fz receptors (Fig. 1a) – has been proposed to act like a stack to accommodate the segments of CRD and Connector on the top of the transmembrane domain^26,29^.

Next, we compute the evolutionary conservation of amino acid sequences of the segments of CRD and Connector for the construction of the phylogenetic tree (Fig. 1b) by the neighbourjoining method^30,31^. Interestingly, the obtained phylogenetic analysis presents the same evolutionary pattern as using either CRD and full-length sequences of Fz receptors^2,24^. As shown in Fig. 1b, ten homologs of Fz receptors divide into five subtypes: (1) Fz1/2/7, (2) Fz5/8, (3) Fz4, (4) Fz9/10, and (5) Fz3/6, consistent with the current understanding of binding specificity of Wnt and Fz pairs and functional redundant and complement of Fz receptors in genetic studies.

### Structures of Fz4 and Fz7 in complex with the palmitoleoylated Wnt7A peptides

One of the long-standing questions in Wnt signalling is how Fz receptors progress the receptor oligomerization to the Wnt signalosome formation through the acylated Wnt binding. To shed light on this question, we expressed two distinct evolutionary branches of Fz receptors – human Fz4 containing the segments CRD and Connector (Fz4_CRD-Connector_) and human Fz7 with the segment of CRD (Fz7_CRD_) in mammalian cells. We co-crystalized Fz4_CRD-Connector_ and Fz7_CRD_ proteins with PAM peptides – palmitoleoylated disulfide-bounded Wnt7A peptides (Cys202-Cys209), which also has been used for structural studies of Notum, a carboxylesterase Wnt specific deacylase^12^, and Glypican, a modulator for Wnt mobility^11^. Next, we determined the crystal structures of Fz4_CRD-Connector_ bound to PAM peptide in two crystal forms at 1.8 Å and 2.95 Å resolutions, respectively, and Fz7_CRD_ in complex with PAM peptide at 1.92 Å resolution (Fig. 2, Supplementary Table 1). In general, these structures adopt the canonical Fz_CRD_ fold, which contains four alpha-helices, connecting loops, five disulfide bridges, and a hydrophobic groove for acrylated Wnt binding^9,24,32,33^. Interestingly, two crystal forms of the Fz4_CRD-Connector_:PAM peptide complex adopted a homodimeric arrangement and presented as an open-form (Figs. 2a and 2b), in contrast to a dimeric closed-form of Fz7_CRD_:PAM peptide complex (Fig. 2c). These dimeric arrangements (open-form and closed-form) adopt the enlarged hydrophobic grooves for the ligand binding by meting two hydrophobic grooves of each monomer at the dimer interface but reveal two distinct types of hydrophobic grooves – elongated and curved shapes for dimeric Fz4_CRD-Connector_ and dimeric Fz7_CRD_, respectively (Figs, 3a, 3b, and 3c). Furthermore, either dimeric open-form or closed-form accommodates one molecule of PAM peptides that appear to coordinate with the dimerization to assemble these enlarged hydrophobic grooves (Figs. 2 and 3). Of note is that three distinct conformations of PAM peptides were observed in the structures of Fz4_CRD-Connector_ and Fz7_CRD_, showing the dynamic mobilities of PAM peptides (Fig. 2). Moreover, residues around the dimer interface for the hydrophobic grooves are conserved among Fz receptors and shared by each monomer (Figs. 3d, 4e, and 4f, Supplementary Fig. 1), no matter open-form and closed-form of the dimer arrangements (Fig 3). Collectively, these results suggest that acylated Wnt binding mediates the dimerization of Fz_CRD_ receptors, in agreement with previous studies showing a dimeric open-form of Fz4_CRD_ in complex with PAM^23^ and a closed-form dimer of Fz5_CRD_ in complex with PAM^24^.

**Figure 2.**
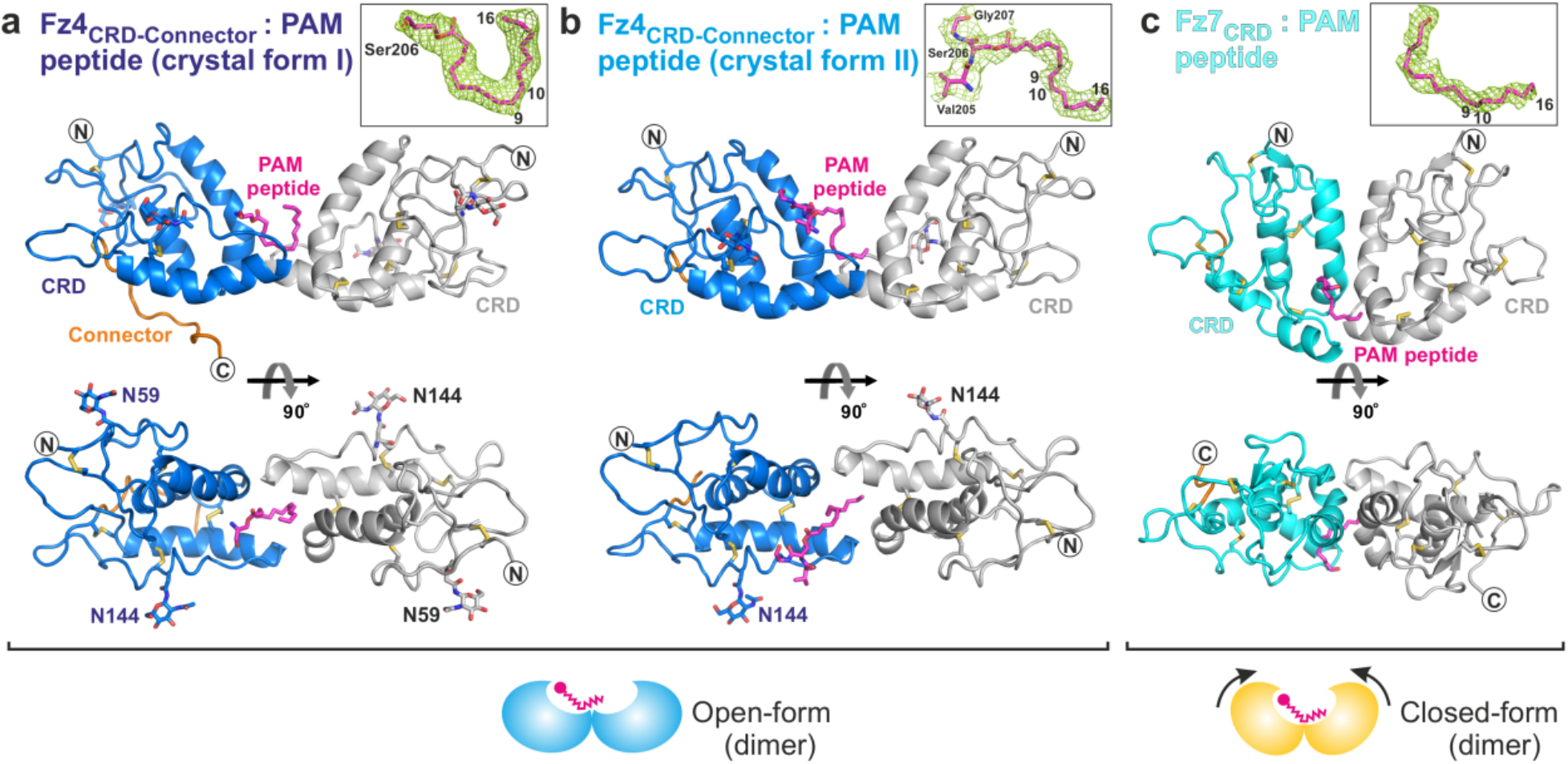
Structural basis of Fz4 and Fz7 in complex with the palmitoleoylated Wnt7a peptide. Ribbon diagrams of Fz4_CRD-Connector_ (CRD, blue and grey; Connector, orange) in complex with PAM peptide (magenta) are shown in an open-form of dimeric assembly – **(a)** crystal form I and **(b)** crystal form II. **(c)** Closed-form of dimeric assembly of Fz7_CRD_ (cyan and grey) in complex with PAM peptide. Insets show the close-up view of the PAM peptide and the 2|*F*_O_|-|*F*_C_| electron density map (green meshes) contoured at 1σ. Asn-linked glycans on N59 and N144 residues, disulfide bonds, and PAM peptides are shown as sticks.

**Figure 3.**
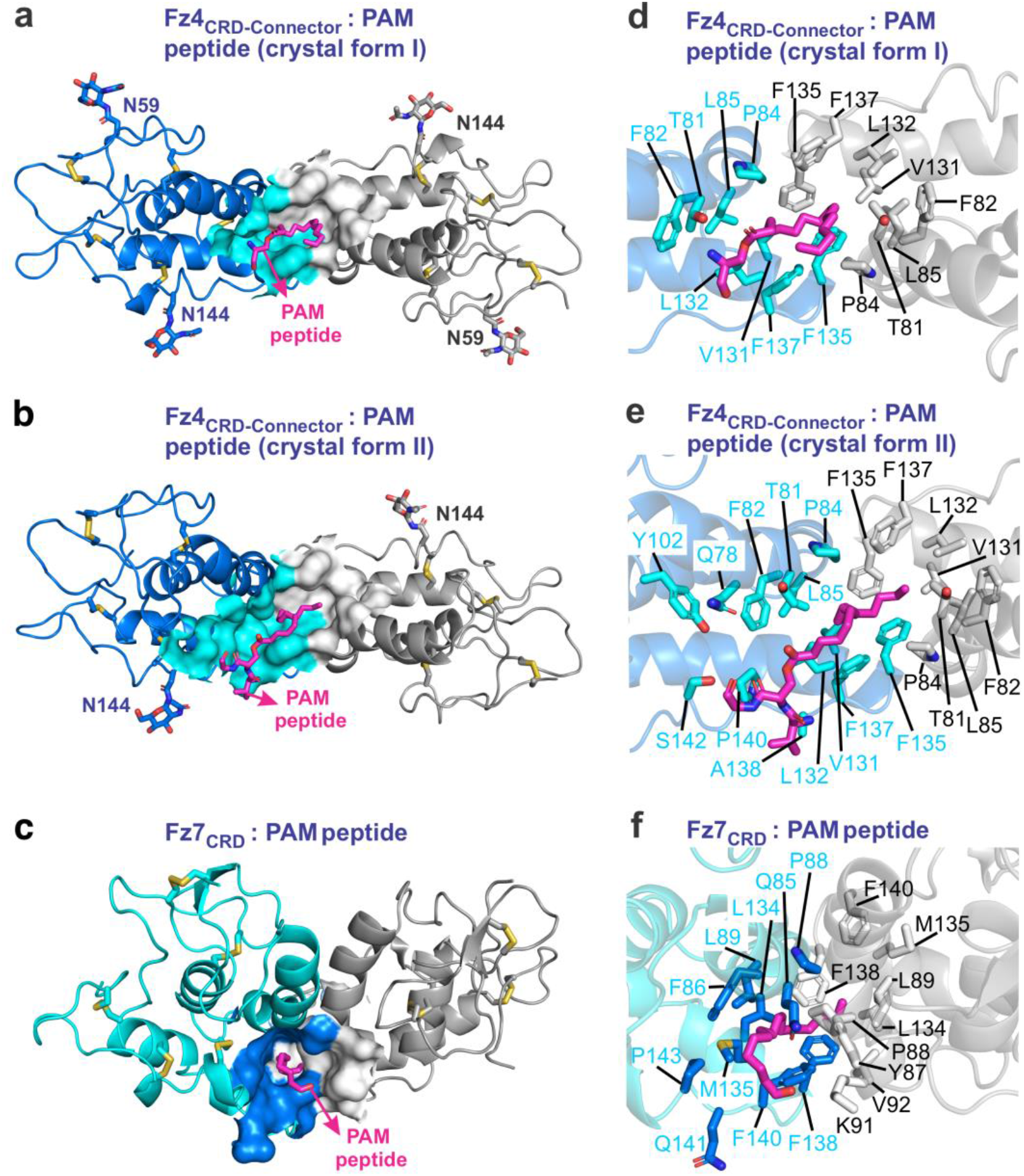
Structural analyses of Fz4 and Fz7 dimer interfaces and the hydrophobic ligand-binding grooves. Hydrophobic ligand volumes (rendered with CASTp using a 1.4 Å probe; shown in cyan and grey for **(a)** and **(b)** and blue and grey for **(c)**) and close-up views of the ligand-binding grooves for the PAM peptide (magenta sticks) binding. Each copy of Fz4 and Fz7 are shown in blue and gray, respectively. **(a)** and **(d)** The crystal form I of Fz4_CRD-Connector_:PAM peptide. **(b)** and **(e)** The crystal form II of Fz4_CRD-Connector_:PAM peptide. **(c)** and **(f)**Fz7_CRD_:PAM peptide.

**Figure 4.**
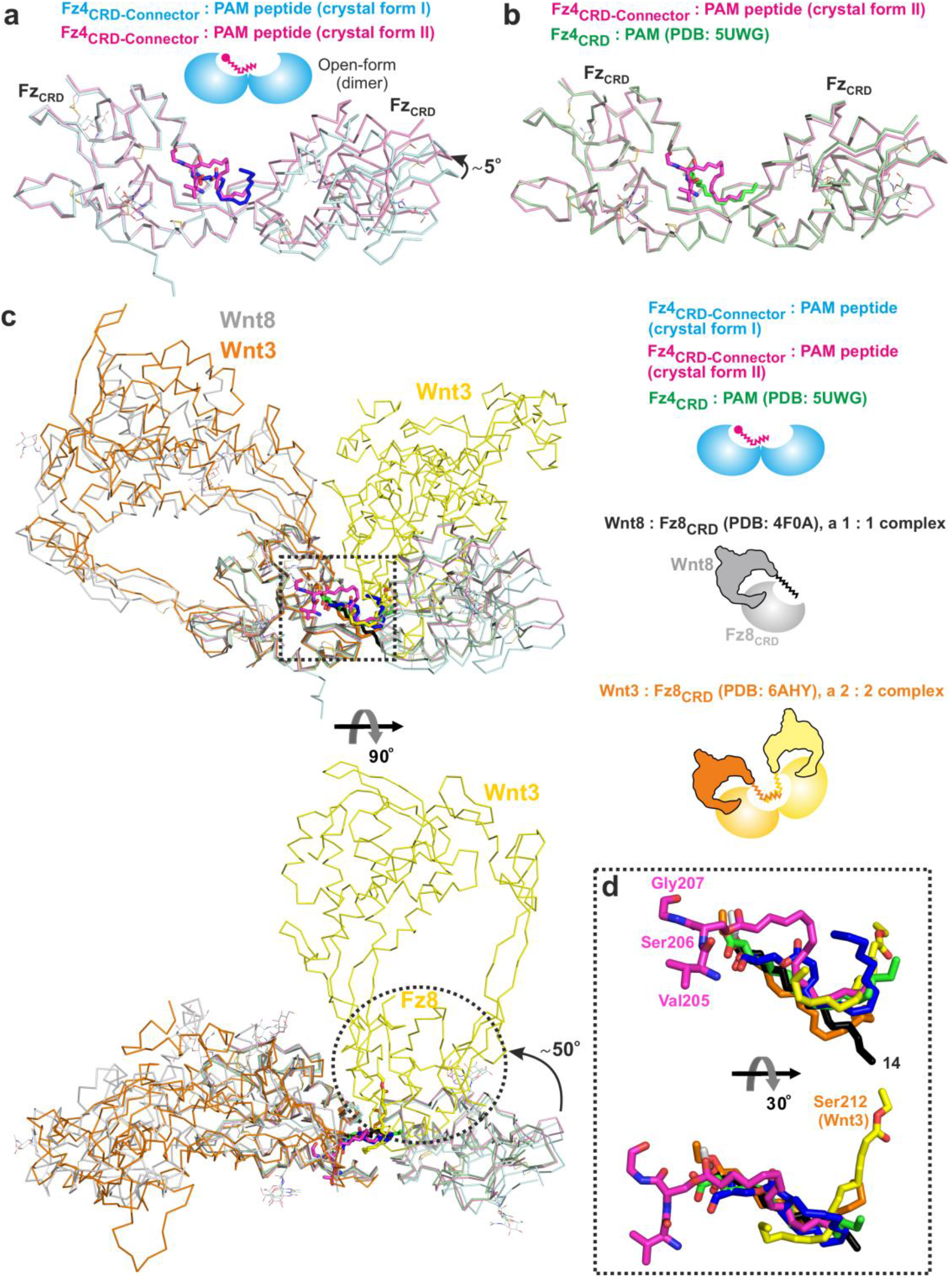
Structural analyses of the dimeric open-form assembly. **(A)** Superposition of crystal form I (cyan) and II (pink) of Fz4_CRD-Connector_:PAM peptide with r.m.s. deviation of 0.93 Å over 226 Cα atoms. PAM peptides are shown as sticks (crystal form I, blue; crystal form II, magenta). **(B)** Superposition of Fz4_CRD-Connector_:PAM peptide (crystal form II) and Fz4_CRD_ in complex with PAM (green; PDB: 5UWG) with r.m.s. deviation of 0.48 Å over 209 Cα atoms. **(C)** Ribbon diagram showing a superposition of Fz4_CRD-Connector_:PAM peptide (crystal form I, cyan; crystal form II, pink), Fz4_CRD_:PAM (green), Wnt8 complexed with Fz8_CRD_ (a 1:1 stoichiometry showing in grey; PDB: 4F0A), and Wnt3 complexed with Fz8_CRD_ (a 2:2 stoichiometry showing in orange and yellow; PDB: 6AHY) and the left inset showing their cartoon models of the protein complex assembly. The ligand-binding pocket is marked with a black dotted square. **(D)** A close-up view of the PAM peptide (blue, Fz4_CRD-Connector_ crystal form I; magenta, Fz4_CRD-Connector_ crystal form II), PAM (green, Fz4_CRD_), and PAM attached on Wnt8 (black) and Wnt3 (organ and yellow) as shown in **(C)**.

### Structural analyses of the dimeric open-form and closed-form arrangement

To dissect the distinct dimeric arrangement for acylated Wnt binding, we firstly superposed two crystal forms of Fz4_CRD-Connector_:PAM peptide, which revealed that (1) r.m.s. deviation of 0.93 Å over 226 Cα atoms, (2) conservation of binding sites on Fz4 for the two distinct conformations of PAM peptides, and (3) approximately 5° rotation towards the centre dimer interface from crystal form I to II (Fig. 4a). The overall dimeric arrangement of Fz4_CRD-Connector_:PAM peptide (crystal form II) resembles a reported structure of Fz4_CRD_ in complex with PAM^23^ with r.m.s. deviation of 0.48 Å over 209 Cα atoms (Fig. 4b). The binding sites for PAM peptide and PAM are conserved. As shown in Fig. 4c, we superposed these structures of Fz4_CRD-Connector_ and Fz4_CRD_ with the structures of Wnt8 bound to Fz8_CRD_ in a 1:1 stoichiometry^8^ and Wnt3 bound to Fz8_CRD_ in a 2:2 stoichiometry^25^. According to the structural analyses (Fig. 4c), we note that (1) the superposition of Fz4 dimeric open-form and Wnt8 in complex with Fz8_CRD_ supports a 1:2 complex of Wnt to Fz, in agreement with previous studies^23,24^, (2) the superposition of Fz4 dimeric open-forms and Wnt3 in complex with Fz8_CRD_ reveals an approximately 50° rotation of the second copy of Fz moving toward the second copy of Wnt3 for binding, which appears to suggest a transition state of Wnt to Fz from a 1:2 stoichiometry to a 2:2 stoichiometry, and (3) the hydrophobic grooves at the dimer interfaces present the plasticity to accommodate divergent conformational difference of ligands, including PAM, PAM peptide, and PAM on Wnt (Fig. 4d).

Next, we superposed the structure of Fz7_CRD_:PAM peptide with two reported closed-forms of Fz_CRD_ structures^24^ – (1) Fz7_CRD_ in complex with a C24 fatty acid containing 24 carbons which revealed r.m.s. deviation of 1.05 Å over 212 Cα atoms (Fig. 5a), and (2) Fz5_CRD_ in complex with PAM which showed r.m.s. deviation of 1.28 Å over 214 Cα atoms (Fig. 5b). It is noteworthy that a highly similar closed-form of dimer was observed among these three structures (Figs. 5a and 5b) and the conserved dimer interface for ligand binding, although the PAM in Fz5_CRD_ structure moves deeper into another copy of Fz5_CRD_ (Fig. 5b). Interesingly, the superposition of these structures of dimeric closed-forms with the structure of Wnt3 in complex with Fz8_CRD_ strongly support a 2:2 stoichiometry of the Wnt and Fz pairs (Fig. 5c) and the conformations of ligands (PAM, C24, PAM peptide, and PAM on Wnt) are more convergent in these closed-forms (Fig. 5d) than open-forms (Fig. 4d). As shown in Fig. 6a, we also found that the second copy of Fz_CRD_ in the closed-form rotates approximately 50° toward the ligand-binding site from the openform, consistent with the superimposition of Fz4 dimeric open-forms and a 2:2 complex structure of Wnt3 and Fz8_CRD_ (Fig. 4c). Taken together, these analyses allow us to propose a model of how Fz recognizes the acylated Wnt in a 1:1 stoichiometry initially and assembles as a 2:2 stoichiometry from the dimeric arrangement of open-form and closed-form for Wnt signalling activation (Fig. 6b; see Discussion section in detail).

**Figure 5.**
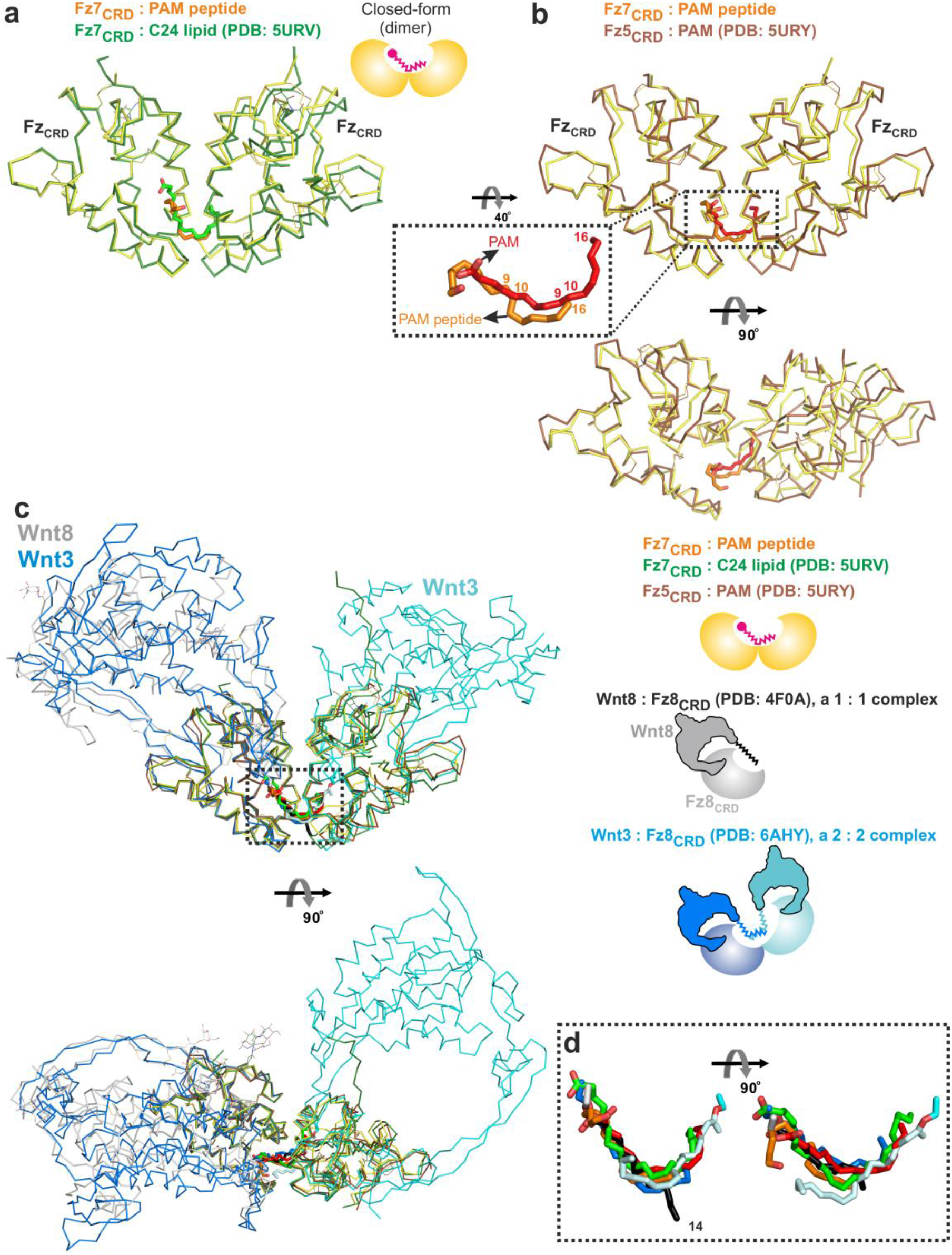
Structural analyses of the dimeric closed-form assembly. **(A)** Superposition of Fz7_CRD_:PAM peptide (yellow) and Fz7_CRD_ in complex with C24 lipid (green; PDB: 5URV) with r.m.s. deviation of 1.05 Å over 212 Cα atoms. **(B)** Superposition of Fz7_CRD_:PAM peptide (yellow) and Fz5_CRD_ in complex with PAM (brown; PDB: 5URY) with r.m.s. deviation of 1.28 Å over 214 Cα atoms and the ligand-binding pocket marked with a black dotted square. The inset shows a close-up view of PAM peptide (orange) from Fz7_CRD_ and PAM (red) from Fz5_CRD_. **(C)** Ribbon diagram showing a superposition of Fz7_CRD_:PAM peptide (yellow), Fz7_CRD_:C24 lipid (green), Fz5_CRD_:PAM (brown), Wnt8 complexed with Fz8_CRD_ (a 1:1 stoichiometry showing in grey; PDB: 4F0A), and Wnt3 complexed with Fz8_CRD_ (a 2:2 stoichiometry showing in blue and cyan; PDB: 6AHY) and the left inset showing their cartoon models of the protein complex assembly. The ligand-binding pocket is marked with a black dotted square. **(D)** A close-up view of the PAM peptide (orange, Fz7_CRD_), C24 lipid (green, Fz7_CRD_), PAM (red, Fz5_CRD_), and PAM attached on Wnt8 (black) and Wnt3 (blue and cyan) as shown in **(C)**.

**Figure 6.**
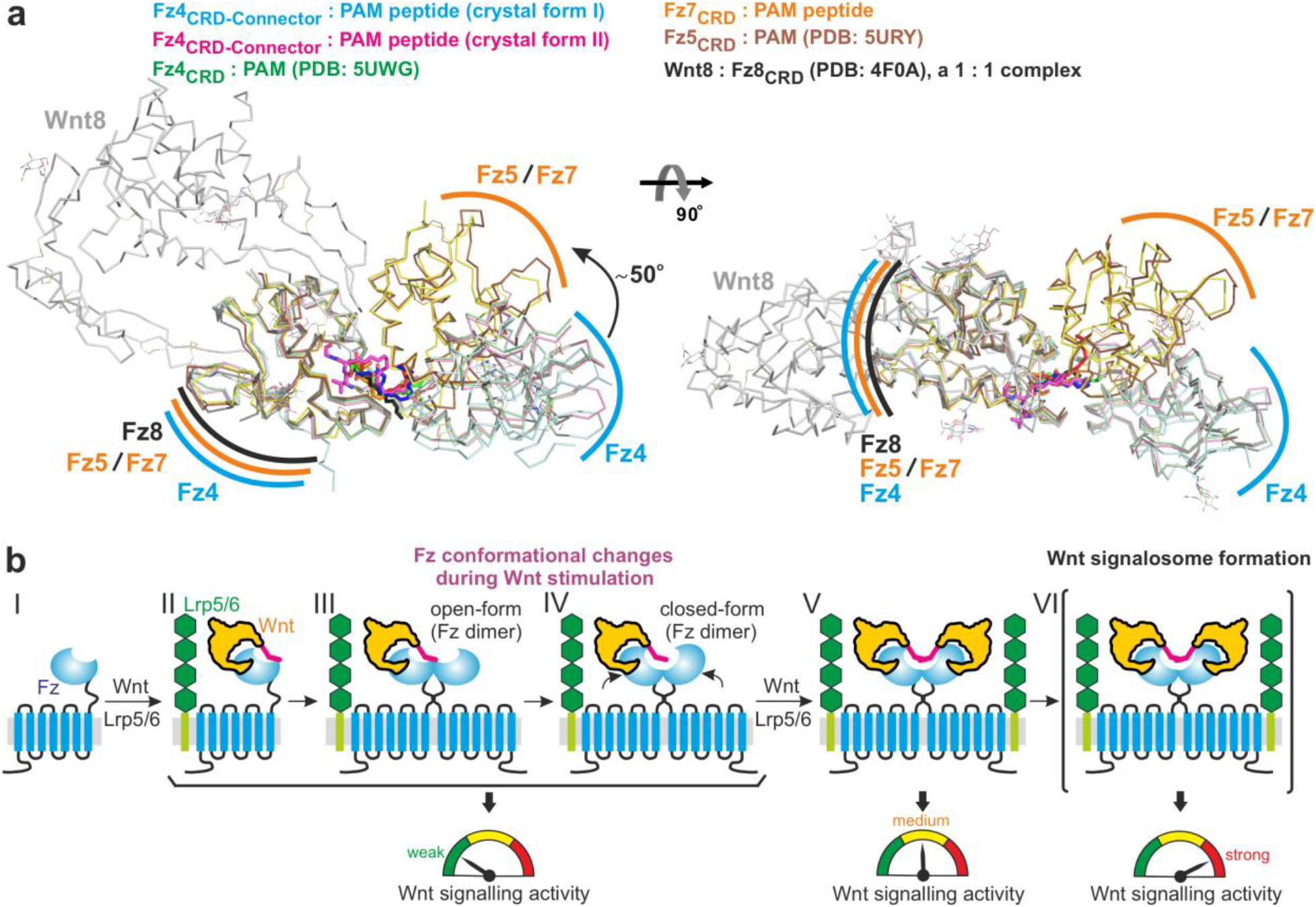
Structural comparison of the dimeric open-form and closed-form assemblies and proposed model of the assembly of Fz receptors in Wnt signalling activation. **(A)** Ribbon diagram showing a superposition of Fz4_CRD-Connector_:PAM peptide (crystal form I, cyan; crystal form II, pink), Fz4_CRD_:PAM (green), Fz7_CRD_:PAM peptide (yellow),, Fz5_CRD_:PAM (brown), and Wnt8 complexed with Fz8_CRD_ (a 1:1 stoichiometry showing in grey). Around a 50°rotation of one copy of Fz molecules the open-form toward closed-form. **(B)** Cartoon model showing that (1) a 1:1 stoichiometry of Wnt and Fz complex with Lrp5/6 triggers the weak Wnt signalling activity, (2) the further Fz dimer assembles from the open-form toward the closed-form, (3) a 2:2 stoichiometry of Wnt and Fz complex occurred stimulates the medium strength of Wnt signalling, and (4) signalosome formation assembled by intracellular Dishevelled oligomerization induces the strong Wnt signalling activity.

### The structure Fz4 Connector and its function in ligand binding

As shown in Fig. 2a, we observed the structure of the Connector segment in crystal form I of Fz4_CRD-Connector_:PAM peptide, which was supported by an electron density (Fig. 7). Interestingly, the segment of Fz4_Connector_ is stacked along helices C and D of Fz4_CRD_ through the length of Fz_CRD_ segment and has extensive hydrophobic and hydrophilic interactions between two segments (Fig. 7, Supplementary Table 2). Specifically, M159 on Fz4_Connector_ shields the hydrophobic pocket, constituted by three residues (Y58, M105, and I114) which were identified in Familial exudative vitreoretinopathy (FEVR) disease (Fig. 7). Of note is that FEVR is a hereditary disorder charactered by abnormal development of the retinal vasculature, often leading to retinal detachment and vision loss^34^. Furthermore, mutations on these residues of (Y58, M105, and I114) have been found to impair the signalling activities^35–37^. Interestingly, a disease-associated residue P168^38^ on the segment of Connector appears to interact with R127^39^, a disease-associated residue on helix D of the CRD segment (Fig. 7, Supplementary Table 2). The roles of R127 and P168 in the signalling activity and disease development will need to determine. Furthermore, our structural analyses (Fig. 7) found that the interactions between Fz4 segments of CRD and Connector orient the Connector toward a specific direction, very likely to the plasma membrane. This orientation of Connector appears to not affect Fz binding to Wnt; however, the substitutions of Connector residues may affect the interaction of the C-terminal lobe of Wnt with Fz (Supplementary Fig. 2). Interestingly, the interactions between Connector and CRD orient the Connector toward the transmembrane domains, in agreement with the observation in SMO full-length structures^26–28^, although the mobility orientation of connectors is different between Fz4 and SMO (Fig. 8a). Moreover, the PAM binding site on Fz4 is located at the position where the cholesterol bound to the CRD of SMO (Fig. 8b). It is noteworthy that cholesterol did not mediate the dimerization of SMO.

**Figure 7.**
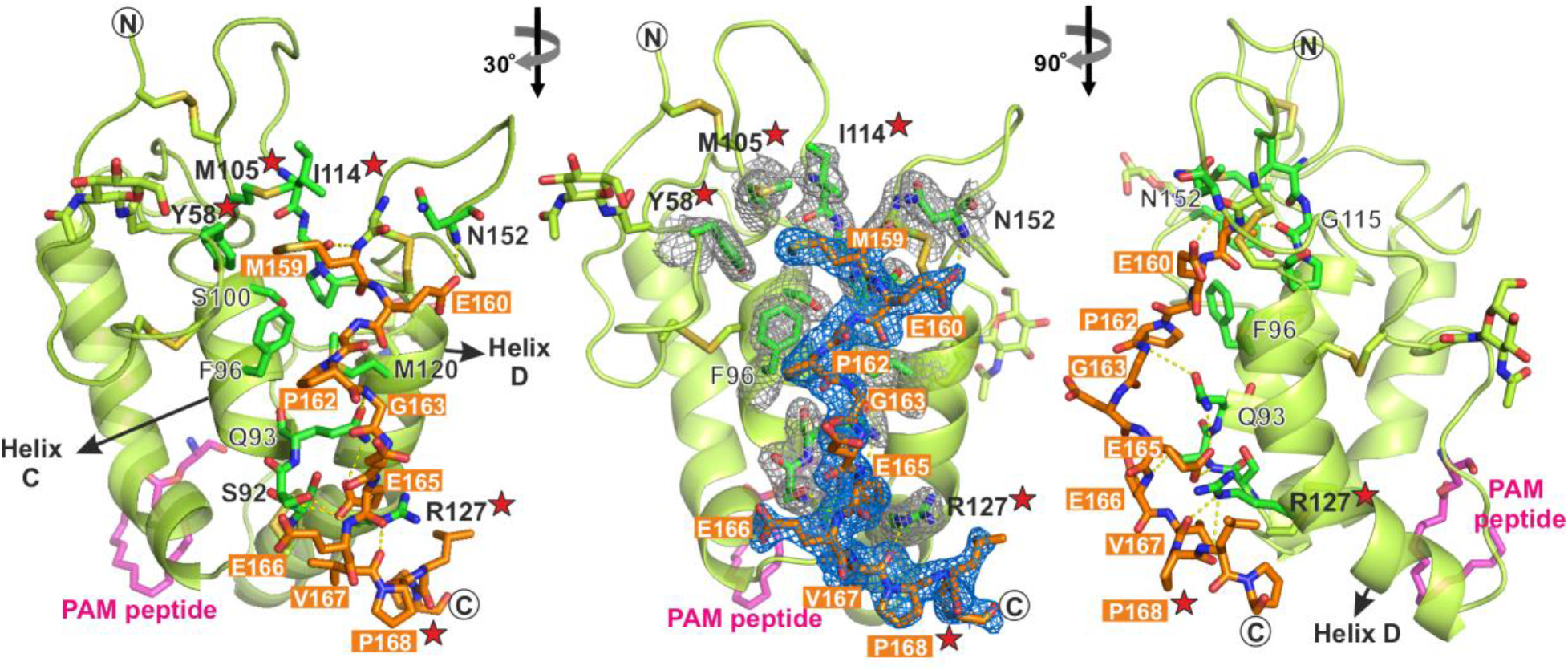
The Connector segment of Fz4 and its interactions with the CRD segment of Fz4. The connector region is highlighted as orange sticks and the CRD is presented as a cartoon representation in green colour. N-linked glycans, disulfide bonds, and PAM peptides are shown as sticks. Circles denote the N- and C-termini. The residues associated with disease mutations are marked by red stars (Y58, M105, I114, R127, and P168). The 2|*F*_O_|-|*F*_C_| electron density maps contoured at 1σ are shown as blue meshes for the connector and grey meshes for residues on CRD interacting with the connector, respectively. Dotted yellow lines denote hydrogen bonds. Table S2 for the detailed interaction networks.

**Figure 8.**
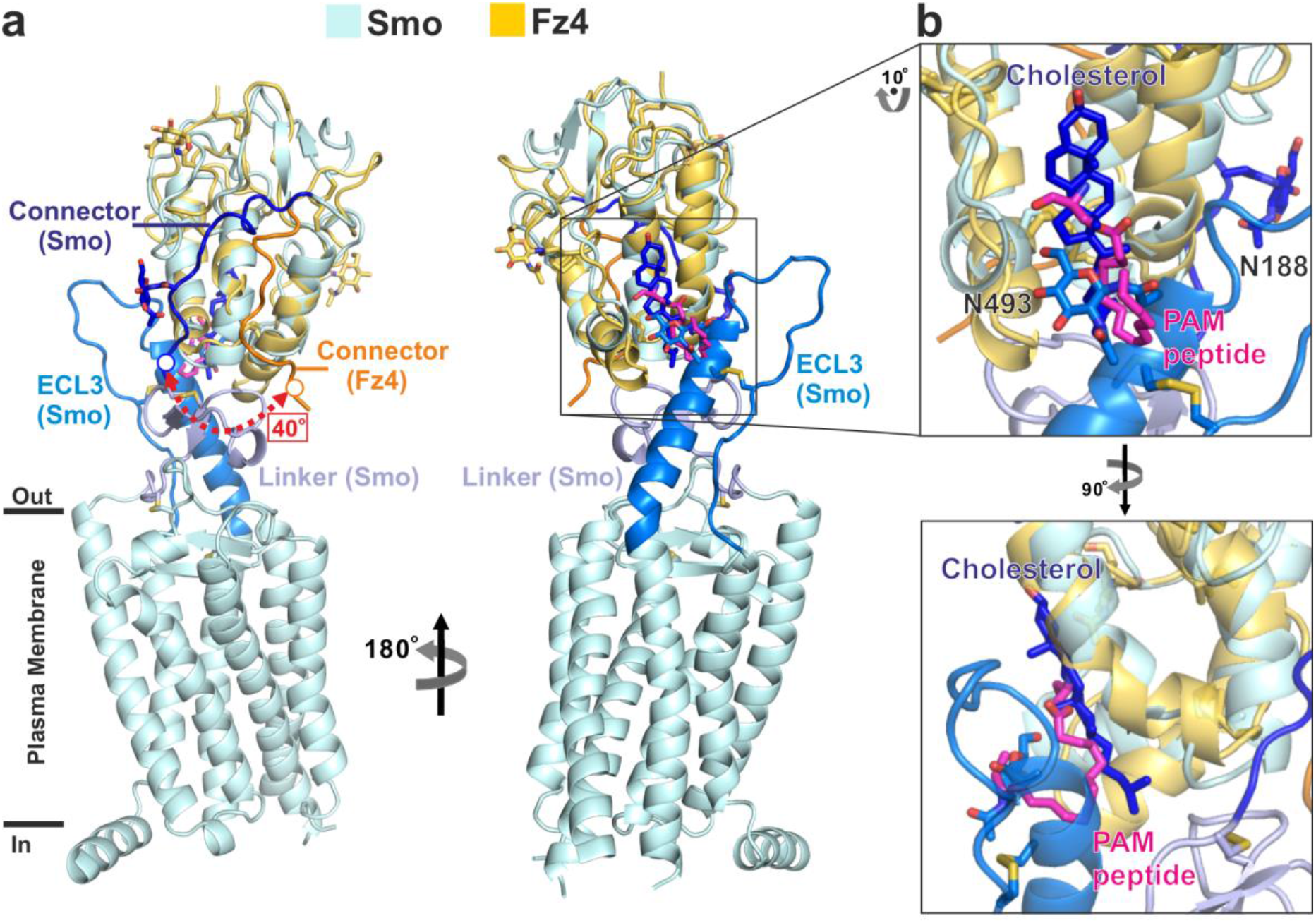
Structural comparison of the Connector segments of Fz4 and Smoothened (Smo) and their ligand-binding sites. **(A)** Superposition of Fz4_CRD-Connector_:PAM peptide (yellow) and Smo in complex with cholesterol (cyan; PDB: 5L7D) shows r.m.s. deviation of 2.79 Å over 90 Cα atoms. The connector regions of Smo (blue) and Fz4 (orange) have around a 40°orientation difference. The ligand-binding pocket is marked with a black square. **(B)** A close-up view of the ligand-binding pocket as shown in **(A)**. The PAM peptide (magenta) of Fz4_CRD-Connector_ and cholesterol (blue) are shown as sticks.

Fz4 also has been well known to mediate β-catenin signalling through the interaction with Norrin (also called Norrie Disease Protein), a secreted cystine-knot like growth factor that mimics Wnt functionally^32,35,40^. To explore the role of Fz4_Connector_, we superposed the structures of Fz4_CRD_-Connector and a reported complex structure of Norrin bound to Fz4_CRD_^32^. We found that the beginning part of Fz4_Connector_ appears to facilitate the interaction with Norrin together with Fz4_CRD_ (Fig. 9a), in agreement with a previous hydrogen deuterium exchange mass spectrometry (HDX-MS) experiment showing that Fz4 full-length bound to Norrin induces the conformational changes of Fz4 around the segment of Connector^41^. Similarly, the beginning part of the Connector segment may also contribute to Wnt and Fz interactions at site 2 of Wnt and Fz_CRD_ complex structures (Supplementary Fig. 2). Of note is that the segment of Connector presents semi-conserved, in contrast to the highly conserved CRD segment (Supplementary Fig. 1).

**Figure 9.**
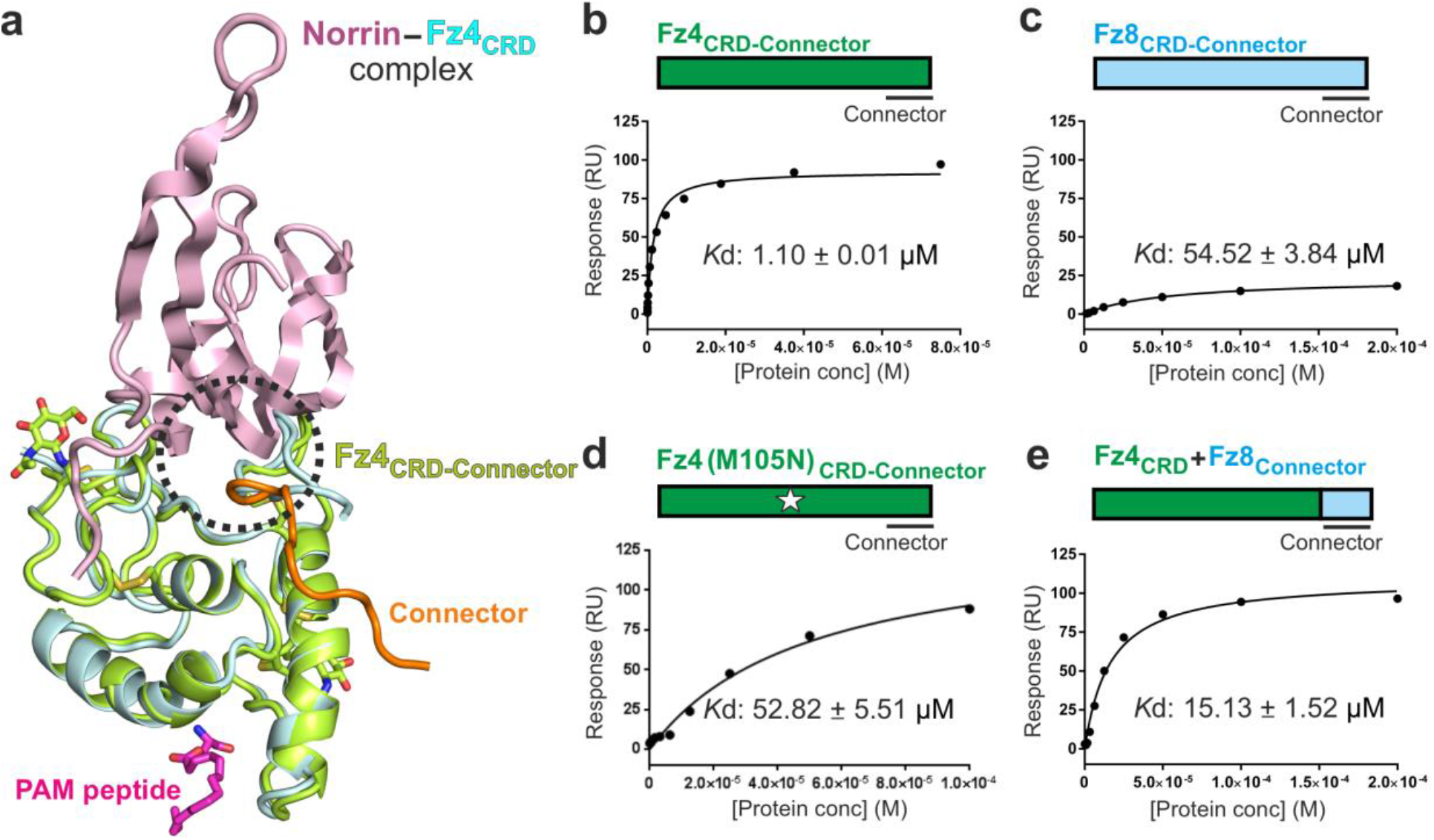
The Connector segment of Fz4 is important for the interaction with the atypical Wnt ligand, Norrin. **(A)** Ribbon representation of Norrin (pink) and Fz4_CRD_ (cyan) complex (PDB: 5BQE), superimposed with one copy of Fz4_CRD-Connector_:PAM peptide (crystal form I; green). The PAM peptide (magenta) and N-linked glycans (green) are shown as sticks. The connector of Fz4_CRD-Connector_ is highlighted in orange. The interaction region between Norrin and the connector of Fz4 is marked with a black dotted circle. **(B)-(E)** SPR results for Fz4_CRD-Connector_, Fz4_CRD-Connector_ (M105N) mutant, Fz8_CRD-Connector_, and the chimaera of Fz4_CRD_+Fz8_Connector_ binding to Norrin.

Next, we hypothesize that Fz4_Connector_ also contributes to Norrin binding. To test the hypothesis, we performed Surface Plasmon Resonance (SPR) binding assay and mutagenesis. The results showed that Fz4_CRD-Connector_ has the *Kd* value of 1 mM affinity to Norrin (Fig. 9b), in contrast to Fz8_CRD-Connector_ having ~55-fold weaker affinity to Norrin (Fig. 9c). Interestingly, mutant M105N of Fz4_CRD-Connector_ (to introduce an N-linked glycosylation site), predicted to impair the interaction between CRD and Connector segments, show ~53-fold weaker affinity to Norrin (Fig. 9d). More importantly, a chimaera mutant, swapping the Connector segment of Fz4 with the corresponding Connector segment of Fz8, reveals ~15-fold weaker affinity for Norrin (Fig. 9e), in agreement with a previous study showing that swapping the sequence between the CRD segment and the first transmembrane domain of Fz4 with the corresponding sequence of Fz5 impaired the Norrin mediated signalling activity and Norrin binding to Fz4^41^. Of note is that Bang et al., define the region between CRD and the first transmembrane domain as Linker^41^. This study divided the region into Connector and Linker domains instead.

## Discussion

Wnt palmitoleoylation is a highly conserved post-translational modification for Wnts, except *Drosophila* WntD, an atypical Wnt without acylation and glycosylation^42–44^. Indeed, previous studies have shown that the PAM on Wnt is a key modification for Wnt binding to Fz and the activation of Wnt signalling^5,8^ as well as the formation of Wnt morphogen gradient^11^. Recently, the discoveries of Notum – a secreted deacylase and negative feedback regulator – have been shown to suppress Wnt signalling through the removal of PAM from Wnts^12,13^. However, Speer et al., suggested that Wnt palmitoleoylation is the independence of Wnt signalling activity, and may facilitate the selectivity of Wnt and Fz pairs instead^45^. Thus, understanding how Fz recognizes acylated Wnt was one of the long-standing questions in Wnt signalling. The structure of Wnt8 in complex with Fz8_CRD_ firstly revealed the structural basis of Wnt and Fz interactions in a 1:1 stoichiometry^8^. The hydrophobic groove of monomeric Fz8_CRD_ for PAM binding showed remarkable sequence conservation among Fz receptors^8,9^. Furthermore, our complex structures of Fz4_CRD-Connector_:PAM peptide (Figs. 2a and 2b) and Fz7_CRD_:PAM peptide (Figs. 2c) revealed that (1) PAM binding mediates the dimerization in a 1:2 stoichiometry of Wnt and Fz (Fig. 3), in agreement with previous studies of Fz4_CRD_ and Fz5_CRD_ structures^23,24^, and (2) each PAM occupies the large continuous hydrophobic groove at the dimer interface (Fig. 3). It is noteworthy that monomeric and dimeric arrangements of Fz_CRD_ molecules share the same conserved residues for their hydrophobic grooves. Moreover, DeBruine *et al*., showed that Wnt5a triggers the oligomerization of Fz4 in the membrane of live cells and observed that the oligomerization can be mediated through Wnt5a binding to Fz4_CRD_ while using the membrane-tethered Fz4_CRD_^23^, which is also consistent with the treatment of Wnt3a induced the oligomerization of Fz4 using bioluminescence resonance energy transfer assays^46^. Petersen *et al*., reported that Wnt5a induced the dissociation and re-association of Fz6 dimer using live cell imaging^47^. Therefore, the dynamic balance between monomeric and oligomeric statuses of Fz receptors appears to be conserved among Fz receptors and to be essential for the function of Fz receptors.

Intriguingly, our structural basis studies (Figs. 4 and 5) revealed that acylated Wnt binding can facilitate the distinct dimeric arrangements of Fz_CRD_ – open-form and closed-form assemblies – in agreement with previous structural studies of Fz4_CRD_ bound to PAM^23^ and Fz5_CRD_ bound to PAM^24^, respectively. More importantly, ours and other studies^23,24^ reported a 1:2 stoichiometry of Wnt and Fz interactions in both dimers of open-form and closed-form, which suggests being a key transition status during the signalling initiation and activation upon Wnt binding. Therefore, we propose a model to explain how the oligomerization of Fz mediates the activation of Wnt signalling (Fig.6b). Firstly, from status I to II, one acylated Wnt recruits one Fz receptor and one Lrp5/6 co-receptor to activate signalling initially. That was supported by the structure of Wnt8 bound to Fz8_CRD_^8^ and surrogate Wnt agonists and synthetic bivalent antibodies showing that this 1:1:1 recruitment is sufficient to activate Wnt signalling^48,49^. Secondly, from status II to IV, the acylated Wnt binding further mediates the dimerization of Fz receptors, particularly through the dimeric arrangement of Fz_CRD_ from an open-form status to a closed-form status. That was supported by a 1:2 stoichiometry of Wnt and Fz interactions in this study and previous reports^23,24^. It is noteworthy that additional studies are required to explore the functional relevance of these observed dimeric arrangements for acylated Wnt binding and signalling. Thirdly, the status V, the formation of a 2:2 stoichiometry of Wnt and Fz engages with more Lrp5/6 co-receptors to achieve higher signalling activity. That was supported by a 2:2 complex structure of Wnt3 bound to Fz8_CRD_ and its computational ternary model of Wnt3:Fz8:Lrp5/6^25^. Of note is that the transition from status IV to V occurred at the cell membrane would need to be investigated. Lastly, the oligomerization of Dishevelled has been well known to trigger the Wnt signalosome formation for the activation of Wnt signalling upon Wnt engaging with Fz receptors and Lrp5/6 co-receptors^50^. Thus, we propose that dimerization of Fz receptors upon Wnt binding together with the recruitment of Lrp5/6 co-receptors and Dishevelled assembly for the Wnt signalosome appears to be a key signalling event to maximum Wnt signalling output. Collectively, this proposed model (Fig.6b) may shed light on the molecular mechanism underlying Wnt signalling activation.

As shown in Fig. 7, Fz4_Connector_ forms strong interactions with Fz4_CRD_ and appears to stabilize Fz4_CRD_ and restrain the orientation of Fz4_CRD_ probably for the ligand binding, similar to the structures of Smoothened^26–28^. In addition, the sequences of Connectors among Fz receptors present semi-conserved (Supplementary Fig. 1). Of note is that Fz receptors have longer lengths of Connectors than Smoothened and the length of Connectors are variable among Fz receptors (Fig. 1). For example, the Connector segment of Fz8 is the longest one, having 79 residues in length. Thus, it is possible that each Connector segment of Fz receptors functions differently in the distinct subtypes of Fz receptors. Interestingly, Ko et al., recently showed that the segments of Connector and Linker contribute to Wnt binding and signalling and the selectivity of Wnt and Fz pairs as well as the oligomerization of Fz receptors ^46^. We also found that several disease-associated mutations (e.g. Y58C^37^, M105V^35^, I114T^51^, R127H^52^, and P168S^38^) in FEVR are located in the interaction networks between Connector and CRD (Fig. 7). Furthermore, our binding assay showed that swapping the segment of Fz4_Connector_ with the corresponding region of Fz8 decreased the binding affinity to Norrin (Fig. 9). Collectively, these lines of evidence suggest that Connector of Fz receptors contributes to ligand binding and Wnt signalling activity. However, the molecular basis of understanding the segments of Connector and Linker among Fz receptors will need to be determined.

This study suggests that the segment of Fz4_Connector_ plays an important role in Wnt/Norrin binding and signalling. Because the sequences of Connectors are unique among each subtype of Fz receptors (Fig. 1, Supplementary Fig. 1), it implies that the segment of Connector has a great potential to be a target for the development of selective binders to modulate Wnt signalling. Of note is that several binders including antibodies have been developed to target the segment of CRD, for example, Wnt surrogates^48^, synthetic peptides^53^, DARPin-based binders^54^, and anti-Fz_CRD_ antibodies^49,55,56^. However, the binding promiscuities among subtypes of Fz receptors are the major concerns (e.g., unwanted toxicities) of using these binders for therapeutic treatments, as Fz receptors play fundamental roles in embryonic development and adult homeostasis. Interestingly, Gumber et al., recently developed an engineered antibody (F7L6), selectively targeting the Fz7 receptor and the Lrp6 co-receptor and mimicking Wnt for the activation of signalling^57^. F7L6 antibody conjugated to the microtubule-inhibiting drug showed promising results in the regression of ovarian tumour growth^58^. More importantly, the binding epitope of the F7L6 antibody against the Fz7 receptor is located on the Connector of the Fz7 receptor^57^. Thus, the segment of Connector among Fz receptors appears to be an attractive therapeutic target for the development of next-generation binders to achieve the targeting specificity for these diseases caused by dysfunction of Wnt signalling.

In summary, our structural results provide new insights into the PAM on Wnt recognized by the Fz receptor. We revealed that (1) PAM on Wnt could mediate the dimerization of Fz_CRD_ molecules through binding to the hydrophobic groove at the dimer interface, (2) a 1:2 stoichiometry of Wnt and Fz pairs, (3) the distinct dimeric arrangements (open-form and closed-form), and (4) the conformational flexibilities of PAM peptides. Indeed, these studies raise several interesting questions required to further investigation. For example, is the dimerization of Fz_CRD_ molecules initially driven by PAM on Wnt or dose the pre-formed dimer of Fz_CRD_ molecules occur first? What is the dynamic equilibrium of dimeric open-form and closed-form of Fz receptors at the cell membrane? What is the relative ratio among Fz receptor alone, Wnt and Fz receptor complex in 1:2 and 2:2 stoichiometries assembled at the cell membrane? Lastly, we provide structural and functional insights into the segment of Fz4_Connector_. The findings may greatly promote the development of therapeutic strategies to target the subtype of Fz receptors specifically by focusing on the Connector of Fz receptors.

## Methods

### Plasmid design and construction

The coding segments of human Fz4 (UniProt code: Q9ULV1; residues 42-179), human Fz7 (UniProt code: O75084; residues 33-173), and mouse Fz8 (UniProt code: Q61091; residues 30-170) were cloned into the pHLsec-mVenus-12H vector^32,59^. For protein-protein interaction experiments, the construct of Norrin was described previously^32^. For the chimaera of Fz4_CRD_+Fz8_Connector_, coding segments of human Fz4 (residues 42-151) with mouse Fz8 (residues 142-170) cloned into the pHLsec-mVenus-12H vector^32,59^. Human Fz4 (M105N) mutant was generated using two-step overlapping PCR experiments. All constructs were confirmed by DNA sequencing.

### Protein expression and purification

HEK293T (ATCC CRL-11268) cells were maintained in a humidified 37° C incubator with 5% CO_2_ in Dulbecco’s Modified Eagle Medium (DMEM, MilliporeSigma) supplemented with 2 mM L-Glutamine (L-Glu, Gibco), 0.1 mM non-essential amino acids (NEAA, Gibco) and 10% [v/v] Fetal Bovine Serum (FBS, Gibco). The FBS concentration was lowered to 2% [v/v] after transfection with the DNA using polyethylenimine (MilliporeSigma 408727) as described previously^60^. Expression constructs were transfected in HEK293T cells in the presence of 4 mM valproic acid^32^. For protein crystallization, media were supplemented with 5 μM of the class I α-mannosidase inhibitor, kifunensine^61^, after transfection. Conditioned media were harvested five days post-transfection and supplemented with 10 mM HEPES, pH 7.5, and 5 mM imidazole. The His-tagged sample was purified by immobilized metal affinity chromatography (IMAC) using Ni Sepharose Excel resin (GE Healthcare Life Sciences). The IMAC eluted sample was dialyzed and treated with His-tagged Human Rhinovirus 3C protease as described previously^32^. The treated sample was further purified by IMAC and was subjected to size-exclusion chromatography (SEC) using HiLoad Superdex 200 pg (GE Healthcare Life Sciences) in 10 mM HEPES, pH 7.5, 0.15 M NaCl.

### Crystallization and data collection

Purified Fz4_CRD-Connector_ and Fz7_CRD_ proteins were incubated with a palmitoleoylated peptide (Wnt7a residues 202-209; Eurogentec)^12^ at a molar ratio of 1:1.25 overnight at 4°C in a buffer containing 10 mM HEPES, pH 7.5 and 50 mM NaCl. Concentrated Fz4_CRD-Connector_ (45 mg/ml) and Fz7_CRD_ (16 mg/ml) proteins were subjected to setting up Crystallization trials in the presence of 0.5%*Flavobacterium meningosepticum* endoglycosidase-F1 (Endo-F1) prepared as described previously^32^ for *in situ* deglycosylation^62^. Using a Cartesian Technologies dispensing instrument^63^, the drop vapour diffusion crystallization trials were set up in 96-well Greiner plates by mixing a 100 nl protein solution with a 100 nl reservoir. crystallization plates were stored and imaged via the ROCK IMAGER 1000 at 294 K. Fz4_CRD-Connector_ crystal-form I crystallized in 1.6 M ammonium sulphate, 0.1 M NaCl, 0.1 M HEPES, pH 7.5. Fz4_CRD-Connector_ crystal-form II crystallized in 1.6 M ammonium sulphate, 0.1 M Citrate, pH 4.0. Fz7_CRD_ crystallized in 1.5 M ammonium phosphate, 0.1 M Tris, pH 8.5. For cryoprotection, crystals were transferred into a reservoir solution supplemented with 25% glycerol for crystal-form I of Fz4_CRD-Connector_ and Fz7_CRD_ and with 30% glycerol for crystal-form II of Fz4_CRD-Connector_ and subsequently cryocooled in liquid nitrogen. X-ray diffraction data were collected at 100 °K Diamond Light Source (Oxfordshire, UK) at beamlines I24. Diffraction data were indexed, integrated, and scaled using the XIA2 system^64^, coupled with DIALS^65,66^ and POINTLESS^67^.

### Structure determination, refinement, and analysis

The structures of Fz4_CRD-Connector_:PAM peptide and Fz7_CRD_:PAM peptide were determined by molecular replacement using PHASER^68^ and the Fz4_CRD_ structure (PDB ID 5BQC) which was used as a template to obtain the initial phases. The models were completed by manual building in COOT^69^ and refinement was performed using PHENIX Refine^70^ and REFMAC5^71^ with translation-libration-screw parameterization. MOLPROBITY^72^ was used to validate the models. The crystallographic statistics are listed in Supplementary table 1. Structure-based multiple sequence alignment was performed using Clustal Omega^73^. Structure superposition was performed using the alignment algorithm in the PyMOL Molecular Graphic System (Version 2.2, Schrödinger, LLC). The interior volume of the pocket was calculated using CASTp^74^ with a 1.4 Å radius probe. Schematic figures and other illustrations were prepared using GraphPad Prism (GraphPad Software, LLC) and Corel Draw (Corel Corporation).

### Surface plasmon resonance equilibrium binding studies

SPR experiments were performed using a Biacore T200 machine (GE Healthcare Life Sciences) at 25°C in 10 mM HEPES, pH 7.5, 0.15 M NaCl, 0.005% Tween20. For *in vivo* biotinylation, the plasmid DNA mix of Norrin vector and pHLsec-BirA-ER^32^ into a 3 to 1 ratio transfected in HEK293T cells in the presence of 0.1 mM Biotin (MilliporeSigma B4639). The biotinylated Norrin samples were immobilized onto the surface of a CM5 sensor chip (GE Healthcare Life Sciences) on which approximately 8500 resonance units of streptavidin were coupled via primary amines. The signal from SPR flow cells was corrected by subtraction of a blank and reference signal from a mock-coupled flow cell. In all analyses, the experimental trace returned to the baseline line after a regeneration step with 100 mM phosphate pH 3.7, 2 M NaCl, and 1% Tween20. The data were fitted to a 1:1 Langmuir adsorption model (*B*=*B*_max_*C*/(*K*_d_ + *C*), where *B* is the amount of bound analyte and *C* is the concentration of analyte in the sample) for the calculation of dissociation constant (*K*_d_) values using Biacore Evaluation software (GE Healthcare Life Sciences). Data points correspond to the average from two independent dilution series.

## Acknowledgements

We thank Diamond Light Source for beamtime and the staff at beamline I24 for assistance with data collection. Margaret Jones and Tom Walter for technical support. Weixian Lu and Yuguang Zhao for help with tissue culture. XXX for comments on the manuscript. This work was supported by grants to EYJ from XXX. The WTCHG is supported by the Wellcome Trust (XXX). TC was supported by the Human Science Frontier Program Organization Fellowship (LT000130/2017-L).

## Author contributions

Conceptualization: TC, EYJ. Methodology: TC, FH. Investigation TC, KH, FH. Data analysis and visualization: TC, FH, EYJ. Funding acquisition: TC, EYJ. Supervision: TC, EYJ. Writing: TC, EYJ.

## Additional information

Coordinates and structure factors have been deposited in the Protein Data Bank (PDB) with an accession code XXXX, XXXX, and XXXX for Fz4_CRD-Connector_:PAM peptide (crystal form I), Fz4_CRD-Connector_:PAM peptide (crystal form II), and Fz7_CRD_:PAM peptide, respectively.

## References

1 Nusse, R. & Clevers, H. Wnt/beta-Catenin Signaling, Disease, and Emerging Therapeutic Modalities. Cell 169, 985–999, doi:10.1016/j.cell.2017.05.016 (2017).

2 Wang, Y., Chang, H., Rattner, A. & Nathans, J. Frizzled Receptors in Development and Disease. Curr Top Dev Biol 117, 113–139, doi:10.1016/bs.ctdb.2015.11.028 (2016).

3 Malinauskas, T. & Jones, E. Y. Extracellular modulators of Wnt signalling. Current opinion in structural biology 29C, 77–84, doi:10.1016/j.sbi.2014.10.003 (2014).

4 Anastas, J. N. & Moon, R. T. WNT signalling pathways as therapeutic targets in cancer. Nature reviews. Cancer 13, 11–26, doi:10.1038/nrc3419 (2013).

5 Willert, K. et al. Wnt proteins are lipid-modified and can act as stem cell growth factors. Nature 423, 448–452, doi:10.1038/nature01611nature01611 [pii] (2003).

6 Takada, R. et al. Monounsaturated fatty acid modification of Wnt protein: its role in Wnt secretion. Dev Cell 11, 791–801, doi:10.1016/j.devcel.2006.10.003 (2006).

7 Hsieh, J. C., Rattner, A., Smallwood, P. M. & Nathans, J. Biochemical characterization of Wnt-frizzled interactions using a soluble, biologically active vertebrate Wnt protein. Proc Natl Acad Sci U S A 96, 3546–3551 (1999).

8 Janda, C. Y., Waghray, D., Levin, A. M., Thomas, C. & Garcia, K. C. Structural basis of Wnt recognition by Frizzled. Science 337, 59–64, doi:10.1126/science.1222879 (2012).

9 Dann, C. E. et al. Insights into Wnt binding and signalling from the structures of two Frizzled cysteine-rich domains. Nature 412, 86–90, (2001).

10 Bhanot, P. et al. A new member of the frizzled family from Drosophila functions as a Wingless receptor. Nature 382, 225–230, doi:10.1038/382225a0 (1996).

11 McGough, I. J. et al. Glypicans shield the Wnt lipid moiety to enable signalling at a distance. Nature 585, 85–90, doi:10.1038/s41586-020-2498-z (2020).

12 Kakugawa, S. et al. Notum deacylates Wnt proteins to suppress signalling activity. Nature 519, 187–192, doi:10.1038/nature14259 (2015).

13 Zhang, X. et al. Notum Is Required for Neural and Head Induction via Wnt Deacylation, Oxidation, and Inactivation. Dev Cell 32, 719–730, doi:10.1016/j.devcel.2015.02.014 (2015).

14 Bilic, J. et al. Wnt induces LRP6 signalosomes and promotes dishevelled-dependent LRP6 phosphorylation. Science 316, 1619–1622, doi:10.1126/science.1137065 (2007).

15 Schwarz-Romond, T. et al. The DIX domain of Dishevelled confers Wnt signaling by dynamic polymerization. Nat Struct Mol Biol 14, 484–492, doi:10.1038/nsmb1247 (2007).

16 Glinka, A. et al. LGR4 and LGR5 are R-spondin receptors mediating Wnt/beta-catenin and Wnt/PCP signalling. EMBO reports 12, 1055–1061, doi:10.1038/embor.2011.175 (2011).

17 Hao, H. X. et al. ZNRF3 promotes Wnt receptor turnover in an R-spondin-sensitive manner. Nature 485, 195–200, doi:10.1038/nature11019 (2012).

18 Koo, B. K. et al. Tumour suppressor RNF43 is a stem-cell E3 ligase that induces endocytosis of Wnt receptors. Nature 488, 665–669, doi:10.1038/nature11308 (2012).

19 de Lau, W., Peng, W. C., Gros, P. & Clevers, H. The R-spondin/Lgr5/Rnf43 module: regulator of Wnt signal strength. Genes Dev 28, 305–316, doi:10.1101/gad.235473.113 (2014).

20 Yu, H., Ye, X., Guo, N. & Nathans, J. Frizzled 2 and frizzled 7 function redundantly in convergent extension and closure of the ventricular septum and palate: evidence for a network of interacting genes. Development 139, 4383–4394, doi:10.1242/dev.083352 (2012).

21 Dijksterhuis, J. P. et al. Systematic mapping of WNT-FZD protein interactions reveals functional selectivity by distinct WNT-FZD pairs. J Biol Chem 290, 6789–6798, doi:10.1074/jbc.M114.612648 (2015).

22 Voloshanenko, O., Gmach, P., Winter, J., Kranz, D. & Boutros, M. Mapping of Wnt-Frizzled interactions by multiplex CRISPR targeting of receptor gene families. FASEB J 31, 4832–4844, doi:10.1096/fj.201700144R (2017).

23 DeBruine, Z. J. et al. Wnt5a promotes Frizzled-4 signalosome assembly by stabilizing cysteine-rich domain dimerization. Genes Dev 31, 916–926, doi:10.1101/gad.298331.117 (2017).

24 Nile, A. H., Mukund, S., Stanger, K., Wang, W. & Hannoush, R. N. Unsaturated fatty acyl recognition by Frizzled receptors mediates dimerization upon Wnt ligand binding. Proc Natl Acad Sci U S A 114, 4147–4152, doi:10.1073/pnas.1618293114 (2017).

25 Hirai, H., Matoba, K., Mihara, E., Arimori, T. & Takagi, J. Crystal structure of a mammalian Wnt-frizzled complex. Nat Struct Mol Biol 26, 372–379, doi:10.1038/s41594-019-0216-z (2019).

26 Byrne, E. F. et al. Structural basis of Smoothened regulation by its extracellular domains. Nature 535, 517–522, doi:10.1038/nature18934 (2016).

27 Zhang, X. et al. Crystal structure of a multi-domain human smoothened receptor in complex with a super stabilizing ligand. Nature communications 8, 15383, doi:10.1038/ncomms15383 (2017).

28 Huang, P. et al. Structural Basis of Smoothened Activation in Hedgehog Signaling. Cell 174, 312–324 e316, doi:10.1016/j.cell.2018.04.029 (2018).

29 Yang, S. et al. Crystal structure of the Frizzled 4 receptor in a ligand-free state. Nature 560, 666–670, doi:10.1038/s41586-018-0447-x (2018).

30 Saitou, N. & Nei, M. The neighbor-joining method: a new method for reconstructing phylogenetic trees. Mol Biol Evol 4, 406–425 (1987).

31 Kumar, S., Stecher, G. & Tamura, K. MEGA7: Molecular Evolutionary Genetics Analysis Version 7.0 for Bigger Datasets. Mol Biol Evol 33, 1870–1874, doi:10.1093/molbev/msw054 (2016).

32 Chang, T. H. et al. Structure and functional properties of Norrin mimic Wnt for signalling with Frizzled4, Lrp5/6, and proteoglycan. eLife 4, e06554, doi:10.7554/eLife.06554 (2015).

33 Chen, P. et al. Structural basis for recognition of frizzled proteins by Clostridium difficile toxin B. Science 360, 664–669, doi:10.1126/science.aar1999 (2018).

34 Criswick, V. G. & Schepens, C. L. Familial exudative vitreoretinopathy. American journal of ophthalmology 68, 578–594 (1969).

35 Xu, Q. et al. Vascular development in the retina and inner ear: control by Norrin and Frizzled-4, a high-affinity ligand-receptor pair. Cell 116, 883–895, doi:S0092867404002168 [pii] (2004).

36 Smallwood, P. M., Williams, J., Xu, Q., Leahy, D. J. & Nathans, J. Mutational analysis of Norrin-Frizzled4 recognition. J Biol Chem 282, 4057–4068, (2007).

37 Zhang, K. et al. An essential role of the cysteine-rich domain of FZD4 in Norrin/Wnt signaling and familial exudative vitreoretinopathy. J Biol Chem 286, 10210–10215 (2011).

38 Dailey, W. A., Gryc, W., Garg, P. G. & Drenser, K. A. Frizzled-4 Variations Associated with Retinopathy and Intrauterine Growth Retardation: A Potential Marker for Prematurity and Retinopathy. Ophthalmology 122, 1917–1923, doi:10.1016/j.ophtha.2015.05.036 (2015).

39 Xiao, H., Tong, Y., Zhu, Y. & Peng, M. Familial Exudative Vitreoretinopathy-Related Disease-Causing Genes and Norrin/beta-catenin Signal Pathway: Structure, Function, and Mutation Spectrums. J Ophthalmol 2019, 5782536, doi:10.1155/2019/5782536 (2019).

40 Shen, G. et al. Structural basis of the Norrin-Frizzled 4 interaction. Cell research, doi:10.1038/cr.2015.92 (2015).

41 Bang, I. et al. Biophysical and functional characterization of Norrin signaling through Frizzled4. Proc Natl Acad Sci U S A 115, 8787–8792, doi:10.1073/pnas.1805901115 (2018).

42 Ganguly, A., Jiang, J. & Ip, Y. T. Drosophila WntD is a target and an inhibitor of the Dorsal/Twist/Snail network in the gastrulating embryo. Development 132, 3419–3429, doi:10.1242/dev.01903 (2005).

43 Gordon, M. D., Dionne, M. S., Schneider, D. S. & Nusse, R. WntD is a feedback inhibitor of Dorsal/NF-kappaB in Drosophila development and immunity. Nature 437, 746–749, doi:10.1038/nature04073 (2005).

44 Chu, M. L. et al. structural Studies of Wnts and identification of an LRP6 binding site. Structure 21, 1235–1242, doi:10.1016/j.str.2013.05.006 (2013).

45 Speer, K. F. et al. Non-acylated Wnts Can Promote Signaling. Cell reports 26, 875–883 e875, doi:10.1016/j.celrep.2018.12.104 (2019).

46 Ko, S. B. et al. Functional role of the Frizzled linker domain in the Wnt signaling pathway. Commun Biol 5, 421, doi:10.1038/s42003-022-03370-4 (2022).

47 Petersen, J. et al. Agonist-induced dimer dissociation as a macromolecular step in G protein-coupled receptor signaling. Nature communications 8, 226, doi:10.1038/s41467-017-00253-9 (2017).

48 Janda, C. Y. et al. Surrogate Wnt agonists that phenocopy canonical Wnt and beta-catenin signalling. Nature 545, 234–237, doi:10.1038/nature22306 (2017).

49 Tao, Y. et al. Tailored tetravalent antibodies potently and specifically activate Wnt/Frizzled pathways in cells, organoids and mice. eLife 8, doi:10.7554/eLife.46134 (2019).

50 Gammons, M. & Bienz, M. Multiprotein complexes governing Wnt signal transduction. Curr Opin Cell Biol 51, 42–49, doi:10.1016/j.ceb.2017.10.008 (2018).

51 Robitaille, J. M. et al. Phenotypic overlap of familial exudative vitreoretinopathy (FEVR) with persistent fetal vasculature (PFV) caused by FZD4 mutations in two distinct pedigrees. Ophthalmic genetics 30, 23–30, doi:10.1080/13816810802464312 (2009).

52 Kondo, H. et al. Genetic variants of FZD4 and LRP5 genes in patients with advanced retinopathy of prematurity. Mol Vis 19, 476–485 (2013).

53 Nile, A. H. et al. A selective peptide inhibitor of Frizzled 7 receptors disrupts intestinal stem cells. Nat Chem Biol 14, 582–590, doi:10.1038/s41589-018-0035-2 (2018).

54 Dang, L. T. et al. Receptor subtype discrimination using extensive shape complementary designed interfaces. Nat Struct Mol Biol 26, 407–414, doi:10.1038/s41594-019-0224-z (2019).

55 Steinhart, Z. et al. Genome-wide CRISPR screens reveal a Wnt-FZD5 signaling circuit as a druggable vulnerability of RNF43-mutant pancreatic tumors. Nat Med 23, 60–68, doi:10.1038/nm.4219 (2017).

56 Raman, S. et al. Structure-guided design fine-tunes pharmacokinetics, tolerability, and antitumor profile of multispecific frizzled antibodies. Proc Natl Acad Sci U S A 116, 6812–6817, doi:10.1073/pnas.1817246116 (2019).

57 Gumber, D. et al. Selective activation of FZD7 promotes mesendodermal differentiation of human pluripotent stem cells. eLife 9, doi:10.7554/eLife.63060 (2020).

58 Do, M. et al. A FZD7-specific Antibody-Drug Conjugate Induces Ovarian Tumor Regression in Preclinical Models. Mol Cancer Ther 21, 113–124, doi:10.1158/1535-7163.MCT-21-0548 (2022).

59 Chang, T. H., Hsieh, F. L., Smallwood, P. M., Gabelli, S. B. & Nathans, J. Structure of the RECK CC domain, an evolutionary anomaly. Proc Natl Acad Sci U S A 117, 15104–15111, doi:10.1073/pnas.2006332117 (2020).

60 Aricescu, A. R., Lu, W. & Jones, E. Y. A time-and cost-efficient system for high-level protein production in mammalian cells. Acta Crystallogr D Biol Crystallogr 62, 1243–1250, doi:10.1107/S0907444906029799 (2006).

61 Chang, V. T. et al. Glycoprotein structural genomics: solving the glycosylation problem. Structure 15, 267–273, (2007).

62 Hsieh, F. L. et al. The structural basis for CD36 binding by the malaria parasite. Nat Commun 7, 12837, doi:10.1038/ncomms12837 (2016).

63 Walter, T. S. et al. A procedure for setting up high-throughput nanolitre crystallization experiments. Crystallization workflow for initial screening, automated storage, imaging and optimization. Acta Crystallogr D Biol Crystallogr 61, 651–657, (2005).

64 Winter, G. xia2: an expert system for macromolecular crystallography data reduction. Journal of Applied Crystallography 43, 186–190, doi:doi:10.1107/S0021889809045701 (2010).

65 Beilsten-Edmands, J. et al. Scaling diffraction data in the DIALS software package: algorithms and new approaches for multi-crystal scaling. Acta Crystallogr D Struct Biol 76, 385–399, doi:10.1107/S2059798320003198 (2020).

66 Winter, G. et al. DIALS: implementation and evaluation of a new integration package. Acta Crystallogr D Struct Biol 74, 85–97, doi:10.1107/S2059798317017235 (2018).

67 Evans, P. Scaling and assessment of data quality. Acta Crystallogr D Biol Crystallogr 62, 72–82, doi:10.1107/S0907444905036693 (2006).

68 McCoy, A. J. et al. Phaser crystallographic software. J Appl Crystallogr 40, 658–674, doi:10.1107/S0021889807021206 (2007).

69 Emsley, P., Lohkamp, B., Scott, W. G. & Cowtan, K. Features and development of Coot. Acta Crystallogr D Biol Crystallogr 66, 486–501, doi:10.1107/S0907444910007493 (2010).

70 Liebschner, D. et al. Macromolecular structure determination using X-rays, neutrons and electrons: recent developments in Phenix. Acta Crystallogr D Struct Biol 75, 861–877, doi:10.1107/S2059798319011471 (2019).

71 Murshudov, G. N. et al. REFMAC5 for the refinement of macromolecular crystal structures. Acta Crystallogr D Biol Crystallogr 67, 355–367, doi:10.1107/S0907444911001314 (2011).

72 Williams, C. J. et al. MolProbity: More and better reference data for improved all-atom structure validation. Protein Sci 27, 293–315, doi:10.1002/pro.3330 (2018).

73 Sievers, F. & Higgins, D. G. Clustal omega. Curr Protoc Bioinformatics 48, 3 13 11–16, doi:10.1002/0471250953.bi0313s48 (2014).

74 Tian, W., Chen, C., Lei, X., Zhao, J. & Liang, J. CASTp 3.0: computed atlas of surface topography of proteins. Nucleic Acids Res 46, W363–W367, doi:10.1093/nar/gky473 (2018).

