## Supplemental data for "Structural insights into Frizzled assembly by acylated Wnt and Frizzled Connector domain"

**Running Title:** Acylated Wnt mediates Frizzled assembly and Frizzled Connector in Norrin/Wnt signalling

**Keywords**: Wnt signalling, Frizzled, palmitoleate, palmitoleoylation, lipid binding protein, cell signalling protein, Norrie disease, Familial exudative vitreoretinopathy, structural biology

### Supplemental data

**Supplementary Table 1.** Data collection and refinement statistics.

| | $Fz4_{CRD\text{-}Connector}$ : Palmitoleoylated Wnt7a peptide (Cys202-Cys209)[a] | $Fz4_{CRD\text{-}Connector}$ : Palmitoleoylated Wnt7a peptide (Cys202-Cys209)[b] | $Fz7_{CRD}$ : Palmitoleoylated Wnt7a peptide (Cys202-Cys209)[a] |
|---|---|---|---|
| Crystal form | I | II | |
| **Data collection** | | | |
| Space group | $P2_12_12_1$ | $C222_1$ | $P2_12_12_1$ |
| Cell dimensions | | | |
| *a*, *b*, *c* (Å) | 71.7, 102.7, 115.2 | 102.1, 111.2, 77.0 | 42.3, 63.2, 104.8 |
| $\alpha, \beta, \gamma$ (°) | 90, 90, 90 | 90, 90, 90 | 90, 90, 90 |
| Wavelength | 0.9996 | 0.9795 | 0.9953 |
| Resolution (Å) | 76.67 - 1.80 (1.83 - 1.80) | 51.05 - 2.95 (3.00 - 2.95) | 54.09 - 1.92 (1.95 - 1.92) |
| $R_{pim}$ (%)[c] | 1.6 (70.0) | 3.9 (69.2) | 9.1 (184.6) |
| $CC_{1/2}$ (%)[d] | 100.0 (48.1) | 99.8 (57.6) | 99.5 (47.2) |
| $I / \sigma I$ | 16.5 (1.0) | 13.4 (1.3) | 6.7 (1.0) |
| Completeness (%) | 100.0 (99.8) | 100.0 (99.2) | 100.0 (96.8) |
| Multiplicity | 13.1 (13.4) | 25.7 (25.2) | 12.4 (10.9) |
| **Refinement** | | | |
| Resolution (Å) | 76.67 - 1.80 (1.83 - 1.80) | 51.05 - 2.95 (3.00 - 2.95) | 54.09 - 1.92 (1.95 - 1.92) |
| No. reflections | 79434 (7512) | 9470 (964) | 22935 (2147) |
| $R_{work}$ / $R_{free}$ | 19.3 / 22.4 | 22.3 / 26.6 | 19.9/ 24.7 |
| No. atoms | | | |
| Protein | 3919 | 1870 | 1851 |
| Ligand/ion | 126 | 55 | 25 |
| Water | 175 | 16 | 162 |
| *B*-factors | | | |
| Protein | 61 | 64 | 35.7 |
| Ligand/ion | 126 | 93 | 54.3 |
| Water | 52 | 65 | 40.3 |
| R.m.s deviations | | | |
| Bond lengths (Å) | 0.019 | 0.01 | 0.006 |
| Bond angles (°) | 1.63 | 1.66 | 0.93 |
| Ramachandran plot | | | |
| Favored (%) | 98.8 | 92.3 | 97.4 |
| Allowed (%) | 1.2 | 7.7 | 2.6 |
| PDB code | | | |

[a]The X-ray diffraction data were collected from a single crystal.
[b]The X-ray diffraction data were merged from two single crystals.
[c]$R_{pim}$: Precision-indicating merging R-factor.
[d]$CC_{1/2}$: correlation coefficients between random half data sets.
[e]R.m.s deviations: root mean square deviation from ideal geometry.
Values in parentheses are for the highest-resolution shell.

**Supplementary Table 2**. The interactions of the Fz4 segment of Connector with the canonical CRD. Resides associated with disease mutations are marked with asterisks.

| **Fz4 Connector** | | **Fz4 CRD** | | **Interaction** |
|---|---|---|---|---|
| **Residue** | **Group** | **Residue** | **Group** | |
| Met159 | side chain | **Tyr58*** | side chain | Hydrophobic |
| Met159 | side chain | Phe96 | side chain | Hydrophobic |
| Met159 | side chain | Ser100 | side chain | Hydrophobic |
| Met159 | side chain | **Met105*** | side chain | Hydrophobic |
| Met159 | side chain | **I114*** | side chain | Hydrophobic |
| Met159 | side chain | Met120 | side chain | Hydrophobic |
| Met159 | backbone O | G115 | backbone N | Hydrogen bond |
| Glu160 | side chain OE1 | Asn152 | backbone N | Hydrogen bond |
| Pro162 | side chain | Phe96 | side chain | Hydrophobic |
| Pro162 | side chain | Met120 | side chain | Hydrophobic |
| Gly163 | backbone N | Gln93 | side chain OE1 | Hydrogen bond |
| Glu165 | side chain OE1 | Gln93 | side chain NE2 | Hydrogen bond |
| Glu166 | backbone N | Ser92 | side chain OG | Hydrogen bond |
| Val167 | backbone O | **Agr127*** | side chain NE | Hydrogen bond |
| **Pro168*** | backbone O | **Agr127*** | side chain NE | Hydrogen bond |

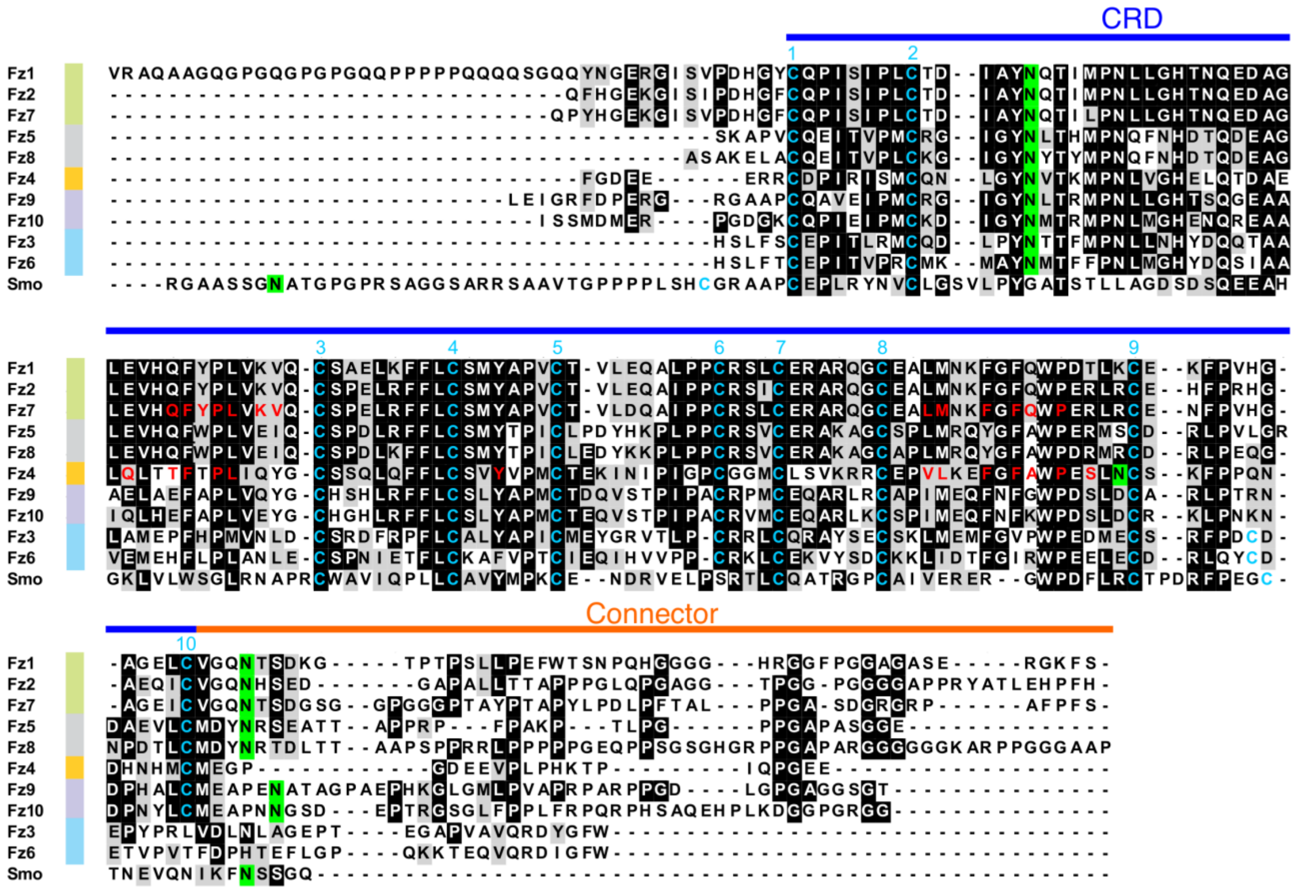

**Supplementary Figure 1 |** Multiple sequence alignment of the segments of CRD and Connector of human Fz receptors and Smoothened (Smo).

CRD, a region between the first and tenth conserved cysteine residues, is drawn in a blue line and the region of Connector is drawn in an orange line. Identical residues are shaded in dark and similar residues in gray. Ten conserved cysteine residues that form five disulphide bonds are highlighted in cyan. Residues of the ligand binding groove for PAM peptide binding are highlighted in red. The asparagine residues of the N-glycosylation sites are highlighted in green. Notably, the residues of 189 to 207 of the Fz8 receptor are omitted from the sequence alignment due to the lowest conservation and longest length of the Fz8Connector.

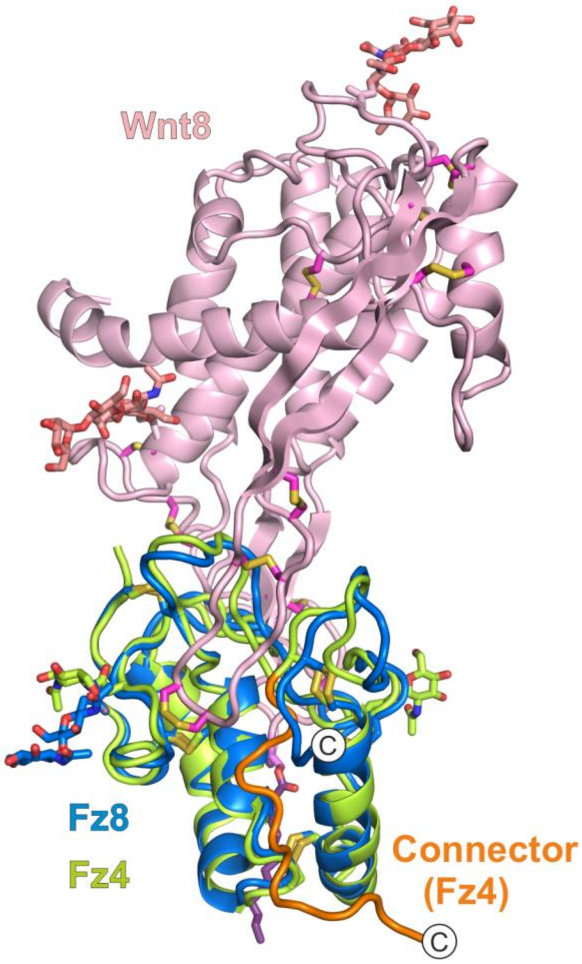


**Supplementary Figure 2 |** Structural comparison of $Fz8_{CRD}$:Wnt8 complex and $Fz4_{CRD\text{-}Connector}$:PAM peptide.

Superposition of Wnt8 (pink) and $Fz8_{CRD}$ (blue) complex (PDB: 4F0A) and $Fz4_{CRD\text{-}Connector}$:PAM peptide (crystal form I; green). The PAM peptide (purple) and N-linked glycans are shown as sticks. The connector of $Fz4_{CRD\text{-}Connector}$ is highlighted in orange. Circles denote the C-termini of $Fz4_{CRD\text{-}Connector}$ and $Fz8_{CRD}$.
